# Human PMS1-dependent non-canonical mismatch repair converges with MBD4 to repair 5-methylcytosine deamination

**DOI:** 10.64898/2026.01.16.698129

**Authors:** Anaïs Le Ven, Sandra Vanhuele, Olivier Ganier, Alexandre Houy, Amanda Kahn, Manuel Rodrigues, Marc-Henri Stern, Raphael Guerois, André Bortolini Silveira

## Abstract

CpG dinucleotides are hotspots for mutagenesis by spontaneous deamination of 5-methylcytosine (5mC) into thymine, resulting in T:G mismatches that can lead to C>T transitions. These mutations are a hallmark of aging and cancer and a major force shaping the evolution of vertebrate genomes. We have previously uncovered MBD4 as the primary base excision repair (BER) glycosylase responsible for 5mC deamination repair. In this study, we employ a cytosine base-editing system, comprising an APOBEC1 deaminase fused to a catalytically dead Cas9, to induce targeted cytosine deamination in native chromatin and track its repair. This approach reveals that MBD4 cooperates with a non-canonical branch of mismatch repair (MMR) to elicit 5mC deamination repair. We demonstrate that MBD4 activity depends on MMR complexes MutSα (MSH2-MSH6) and MutLβ (MLH1-PMS1), but not on post-replicative MMR elicited by the MutLα (MLH1-PMS2) complex. We find that PMS1 loss phenocopies the genome-wide CpG>TpG hypermutated profile associated with MBD4 deficiency, uncovering that 5mC deamination repair may represent one of the primary functions of the MutLβ complex. The mutational landscape of MMR-deficient tumors aligns with our experimental results, showing that replication-independent CpG>TpG mutagenesis partly contributes to the mutational burden of tumors inactivated for *MLH1*, *MSH2* and *MSH6*. Finally, using structural predictions alongside biochemical validation, we show that MBD4 physically interacts with MutLβ in an MLH1-dependent manner, illuminating the structural basis for the convergence of the BER and MMR pathways. Altogether, we uncover a novel function of non-canonical MMR that underscores its interplay with BER in safeguarding genomic integrity against damage to methylated DNA.

## Introduction

DNA methylation is a common epigenetic modification, where a methyl group is added to the 5-position of cytosine residue to form 5-methylcytosine (5mC). In vertebrate genomes, most DNA methylation occurs at CpG dinucleotides (1). CpG dinucleotides are genomic hotspots for mutation, primarily due to the spontaneous deamination of 5mC into thymine. This endogenous DNA damage results in T:G mismatches in double-stranded DNA, leading to C>T transitions if left unrepaired before DNA replication (2). This process is a major source of single-nucleotide variants (SNV), a hallmark of aging and one of the primary contributors to the germline diversity (3,4). Moreover, CpG mutations may play a central role in tumorigenesis, accounting for approximately 30% of the mutational burden in cancer genomes (5,6).

The base excision repair (BER) pathway is involved in the repair of deaminated 5mC through the activity of specialized DNA glycosylases, which recognize T:G mismatches as a substrate and direct excision of the mispaired thymine (7). We uncovered MBD4 (Methyl-CpG Binding Domain Protein 4) as the primary BER glycosylase responsible for 5mC deamination repair in both CpG and non-CpG contexts, with a striking preference for guarding active chromatin and early-replicating DNA (8). Biallelic loss of *MBD4* in cancer is associated with a CpG>TpG hypermutation phenotype and an increased response to immune checkpoint inhibitor therapies (9-12).

DNA mismatch repair (MMR) is a highly conserved biological pathway that plays a key role in maintaining genomic stability. Its primary function is to excise mismatches generated by nucleotide misincorporation during S-phase DNA replication, often referred to as canonical post-replicative MMR. During this process, mismatches are generally recognized by the MutSα (MSH2-MSH6) or MutSβ (MSH2-MSH3) complexes, which then recruit the MutLα complex (MLH1-PMS2). This interaction activates the latent endonuclease activity of MutLα to generate nicks in the discontinuous DNA strand and initiate repair (13). The MutLβ complex (MLH1-PMS1) lacks the motif required for endonuclease activity (14), and its function has remained elusive. Notably, MMR is also active in G_0_/G_1_, often referred to as non-canonical MMR (15). The specialized roles of non-canonical MMR remain poorly understood.

Interestingly, recent evidence suggests that BER and MMR pathways may converge in certain instances of DNA repair (16,17). MBD4 was first identified due to its interaction with MLH1 (18), which enhances efficient recognition of T:G mismatches by MBD4 within methylated CpGs (19). To note, our group and others have shown that germline loss-of-function mutations in either *MBD4* or MMR genes predispose to a partially overlapping spectrum of cancers that includes uveal melanomas and colorectal polyposis (20-22). However, the cooperative roles of BER and MMR pathways in preventing genomic instability remain largely unexplored.

In this study, we define a previously unrecognized pathway for the repair of deaminated 5mC that relies on the coordinated activities of the BER glycosylase MBD4 and a non-canonical MMR axis built around MutSα and MutLβ. Using a targeted Cas9-guided deamination platform, together with genome-wide mutational profiling of engineered cell lines and human tumors, we show that MLH1, PMS1 and MSH6, but not PMS2, are central to limiting CpG>TpG mutagenesis. Our data support a model in which T:G mismatches arising from 5mC deamination are recognized by MutSα, leading to the recruitment of the MBD4-MutLβ complex to initiate repair independently of canonical, post-replicative MMR.

## Methods

### HAP1 isogenic cell models

HAP1 cells were cultured in complete media (IMDM [Gibco, 12440053], bovine fetal serum 10% [BioSera, FB-1003], Penicillin-Streptomycin 100 U/mL [Gibco, 15140122]) at 37 °C in 5% CO_2_. Diploidized clones of HAP1 wild-type or *MBD4* knockout cells (Horizon Discovery, C631 and HZGHC000921c002, respectively) were transfected using jetOPTIMUS (Polyplus, 101000051) with all-in-one CRISPR-Cas9 vectors (pSpCas9(BB)−2A-GFP PX458; Addgene, #48138) with target sequences for *UNG* (AAAGCCCACGGGCACGTTGC), *MLH1* (AAGACAATGGCACCGGGATC), *PMS1* (GTATCCTTAAACCTGACTTA), *PMS2* (ATGCTGTCTTCTAGCACTTC) or *MSH6* (GGAACATTCATCCGCGAGAA). GFP-positive cells were single-cell sorted with SH800S Cell Sorter (Sony Biotechnology) and expanded for approximately 14 days. Clones successfully knocked-out for each gene were identified by western blotting, as described below. The identities of all HAP1 cell models used in this study were confirmed by WGS, as described below. All HAP1 cell models tested negative for mycoplasma with the Eurofins Mycoplasmacheck service, both before and after long-term culturing.

### Cytosine targets selection and bisulfite sequencing

Target cytosines were selected based on pseudo-bulk analysis of whole-genome bisulfite sequencing data of KBM-7 cells (GEO accession number GSE65196) (23), as previously described (8). We opted to target CpGs found within either highly methylated (>99%) or unmethylated (<1%) genomic regions based on average CpG methylation levels across 2 kb genomic bins, and within open-chromatin regions based on chromatin state annotations from ENCODE (dataset ENCFF300WNA). We further selected isolated CpGs not flanked by other cytosines in the same DNA strand within a ±7 bp window from the target CpG. To experimentally confirm the DNA methylation status of the selected target cytosines in HAP1 cells, 500 ng of wild-type HAP1 gDNA was bisulfite-converted with EZ DNA Methylation-Gold Kit (Zymo, ZD5005). PCR on 50 ng bisulfite-converted DNA per reaction was performed with ZymoTaq Polymerase (ZE2001) using the following primers: 5mC target #1 (fwd GGGATTTTATTTTTTTATGTTTTAGTTTT; rev TCATACCTATAATACCAATACTTTAAAAAA), 5mC target #2 (fwd TATTATGTGTTAGATATTGTTAGGAGTT; rev TTTAATTTTTTAACCTAACACACAAAAC), C target #1 (fwd AAAAATGTGGAGAGGGTTAATAAG; rev CTACCTCTCCCTAAACTCTAATAAATTA), C target #2 (fwd AATAATTTAAATGATTTTTTAGTTTTGG; rev ACAAAACCTTTACTATATTTTATTAATCTCT). Agarose gel-purified PCR products were cloned with CloneJET PCR Cloning Kit (ThermoFischer Scientific, K1231), and Sanger sequencing was performed on plasmids of 10 clones per target. High efficiency of bisulfite conversion was confirmed with the Sanger sequencing results.

### dCas9-APOBEC1 system

The CBE variant CMV-YE1-BE3-FNLS-CMV-mCherry (24) (Addgene, #154005) was reengineered with the Q5 Site-Directed Mutagenesis Kit (NEB, E0554S). First, we removed the uracil glycosylase inhibitor (UGI) fused to the Cas9, which was originally intended to inhibit BER and enhance base-editing efficiencies (25). Second, we mutated the D10A nickase Cas9 (nCas9) with the H840 variant to generate a catalytically dead Cas9 (dCas9). Single-guide RNA sequences were cloned into the vector pGL3-U6-sgRNA-EGFP (26) (Addgene, 107721), including for 5mC target #1 in *DNAJC16* (AGATACGAAAGAGATGGTGA), 5mC target #2 in *LRP8* (TGAATCGAGGGGTGGGGAGA), C target #1 in *STK38* (GAGTCGGAGGTTGGGGAAGG), C target #2 in *HNRNPDL* (TAAATCGTGAAATAAAAACA). HAP1 cells were plated in complete media 24h before transfection (1 million cells per well of 6-well plate), and media was replaced with fresh complete media supplemented with 1 μM Palbociclib (CST, 47284S) immediately before co-transfection with 1.4 μg CBE vector and 150 ng of each sgRNA vector per well with jetOPTIMUS (Polyplus, 101000051) at a 1:1 ratio. At 24h post-transfection, media was replaced with fresh complete media supplemented with 1 μM Palbociclib. At 48h post-transfection, cells were harvested, resuspended in FACS buffer containing 1 μM DAPI and live cells were sorted by predefined narrow bands of EGFP and mCherry with a FACSAria Fusion Cell Sorter (BD Biosciences). We fine tuned EGFP and mCherry brightness levels to maximize the quantitative range of base-editing among HAP1 isogenic cells of different genotypes. Genomic DNA was extracted from sorted cells with Quick-DNA Microprep Kit (Zymo, D3021). Custom TaqMan SNP Genotyping Assays (ThermoFischer Scientific, 4331349) were used to perform Droplet Digital PCR (ddPCR) quantitative genotyping using the QX600 ddPCR System (Bio-Rad), separately for each cytosine target. The amount of 10-15 ng DNA was used per reaction with the ddPCR Supermix for Probes no dUTP (Bio-Rad, #1863024). The fractional abundances of DNA strands containing T vs. C in target cytosine positions were used for statistical analysis.

### HAP1 Whole-Genome Sequencing (WGS)

HAP1 cells were split every 3-4 days and kept at low density (< 50% confluency) for approximately 60-90 days following knockout clone selection and initial expansion. A second clonal step was performed by morphology-based single-cell sorting with SH800S Cell Sorter. Expanded subclones had their DNA extracted with DNeasy Blood & Tissue Kit (Qiagen, 69504) and quantified using the Qubit dsDNA BR Assay Kit (Invitrogen, Q32850). 1 μg DNA was used to prepare WGS libraries using the Kapa HyperPrep kit (Roche, 07962363001). Paired-end libraries (2 × 150 bp) were sequenced on a NovaSeq 6000 instrument. WGS coverage depth was set to 25X. Sequencing data QC and adapter trimming on all samples was performed with FastQC v0.11.9 and TrimGalore v0.6.10. Mapping was performed with BWA-MEM v0.7.17 (https://github.com/lh3/bwa) to GRCh38 (Primary assembly + ALT contigs). Duplicates were removed with Sambamba v1.0 (https://github.com/biod/sambamba). All samples had > 90% mapping rate and average fragment size >300 bp. Somatic short variant and insertion/deletion discovery (SNVs + Indels) was performed with GATK4 v4.4.0.0 (https://github.com/broadinstitute/gatk) Mutect2 in ‘tumor’ versus ‘normal’ mode, using sequencing data from HAP1 wild-type parental cells as the normal reference. Variants with a PASS call were selected using the *FilterMutectCalls* function and annotated with VEP v104.3 (https://github.com/Ensembl/ensembl-vep). Variants with less than 4 alternative allele reads or variant allele frequency (VAF) < 0.25 were excluded. Variants shared among subclones of each genotype were further excluded, as they represented variants most likely acquired before knockout first clonal selection step. Estimation of background SNV mutational burden due to culturing was performed with SBS mutational signature analysis with R 4.2.1 package signature.tools.lib v2.4.1 (https://github.com/Nik-Zainal-Group/signature.tools.lib) with the FitMS function in constrained Fit multi-step mode with threshold percent and minCosSimRareSig turned off. The background mutational signature was considered as the average of mutational profile of HAP1 wild-type subclones (two from this study and four from Silveira *et al.* (8)), which was assigned as a common mutational signature. MMR gene knockout signatures previously obtained in human iPSC cells (27) and *MBD4*-deficiency signature in human tumors (SBS96) (28) were assigned as rare mutational signatures. This allowed us to estimate the proportion of the mutational burden in each subclone attributable to background mutagenesis or to mutagenesis associated with the knockout of specific DNA repair genes.

### Human tumors Whole-Genome Sequencing analysis

WGS data from in-house patients, from publicly available *MBD4*-deficient tumors and from the Genomics England [GEL] data release v17 (2023-03-30) were selected, filtered and processed as previously described (8). SBS mutational signature analysis of tumor samples was performed on filtered somatic variants with signature.tools.lib v2.4.1, as previously described (8). MMR-deficient samples were kept for downstream analyses if they showed: (i) any contribution of MMR-deficiency rare signatures SBS6, SBS15, SBS26, SBS44 and SBS97; (ii) no contribution of other rare signatures, including signatures linked to MMR-deficiency in combination with *POLE* or *POLD* dysfunctions; (iii) germline or biallelic somatic loss-of-function mutations in a single of the MMR genes *PMS2*, *MLH1*, *MSH6* or *MSH2*. Replication strand asymmetry analysis was performed with R 4.2.1 package MutationalPatterns v3.8.1 (https://github.com/UMCUGenetics/MutationalPatterns). Replication strand asymmetry of each substitution class (CpG>TpG or other C>T mutations) was defined to be significant if the FDR was <0.01, the absolute value of the log2 ratio was >0.8 and the contribution of the substitution class to the signature was >10%.

#### Recombinant MBD4 overexpression

HAP1 cells stably overexpressing N- or C-terminally FLAG-tagged MBD4 were obtained as previously described (8). An additional synthetic lentiviral vector for overexpression of 3xFLAG N-terminally tagged wild-type *MBD4* (NM_003925.3) followed by IRES and EGFP gene, under the control of the hPGK promoter, was obtained from VectorBuilder. This vector was further modified with the Q5 Site-Directed Mutagenesis Kit (NEB, E0554) to generate the Y417A-F418A mutant of MBD4. Lentiviral particles were generated in HEK-293 and used to transduce HAP1 cells. Cells expressing each construct within a narrow band of EGFP expression were sorted with a SH800S Cell Sorter. Identity of the constructs expressed by each isogenic cell model was confirmed by PCR on genomic DNA followed by Sanger sequencing.

#### Co-immunoprecipitation (co-IP)

For co-IP experiments involving G_1_-cell cycle arrest, HAP1 cells stably overexpressing N- or C-terminally FLAG-tagged MBD4 were cultured in complete media supplemented with 1 μM Palbociclib or vehicle 0.1% DMSO for 17h. For other co-IP experiments, HAP1 cells were cultured in complete media without further treatment. Total cell lysates were obtained from approximately 20 million cells in 300 μL lysis buffer (Tris HCl pH 8, 50 mM; NP-40, 1%; EDTA pH 8, 0.2 mM; 1X protease inhibitor cOmplete EDTA-free [Roche, 11836170001]) at 300 mM NaCl and supplemented with benzonase 250 U/mL, followed by 15 min incubation at room temperature and 15 min at 4°C under agitation. Soluble fractions were obtained by centrifugation at 16,000 g for 15 min at 4 °C and collection of the supernatants. Protein concentrations were measured with the Pierce BCA Protein Assay Kit (Pierce, 23227). Lysates were diluted in lysis buffer to reach final concentrations of 1 μg/μL protein and 150 mM NaCl (input). For co-IP reactions using FLAG-tagged MBD4 as bait, 500 μL diluted lysates and 50 μL pre-washed Pierce ANTI-DYKDDDK magnetic beads (Pierce, A36798) were incubated for 2h at 4°C under agitation. Beads were washed 5 times with 1 mL lysis buffer 150 mM NaCl, and samples were eluted in 2x Laemmli Sample Buffer (Bio-Rad #1610737) with 50 mM DTT for 5 min at 95°C. For co-IP reactions using endogenous proteins as bait, 500 μL diluted lysates were incubated overnight at 4°C under agitation with antibodies against MLH1 (2 μg; CST, 4256S), PMS1 (10 μg; Invitrogen, PA5-35952) or equivalent amount of IgG control (CST, 3900S), followed by 1h incubation at room temperature under agitation with 25 μL pre-washed Pierce Protein A/G Magnetic Beads (Pierce, 88802). Beads were washed 5 times with 1 mL lysis buffer 150 mM NaCl and 0.1% NP-40, samples were eluted in 2x Laemmli Sample Buffer (Bio-Rad #1610737) with 50 mM DTT for 5 min at room temperature, and supernatants were incubated for 5 min at 95°C. Samples were analyzed by western blotting, as described below.

#### Western blotting

For knockout confirmation, HAP1 total cell lysates were extracted in RIPA buffer supplemented with 1X protease inhibitor cOmplete (Roche, 11697498001) and sonicated with Bioruptor Pico (Diagenode) for 10 cycles of 30s on and 30s off at 4 °C. Soluble fractions were obtained by centrifugation at 16,000 g for 15 min at 4 °C and collection of the supernatants. Protein concentrations were measured with the Qubit Protein Broad Range Assay (Invitrogen, A50668). Samples were run in NuPAGE 4 to 12% Bis-Tris gel (Invitrogen, WG1402BOX) or NuPAGE 3 to 8%, Tris-Acetate gel (Invitrogen, WG1602BOX) and transferred to 0.45 µm nitrocellulose membranes. Membranes were blocked with 5% non-fat milk in TBS-T (Tris-Buffered Saline, 0.1% Tween-20), and incubated with primary antibodies in 5% bovine serum albumin (BSA) in TBS-T overnight at 4°C. Antibodies against UNG (1:2000; ProteinTech, 12394-1-AP), MBD4 (1:1000; Abcam, ab224809), FLAG (1:2000; CST, 14793S), MLH1 (1:1000; Sigma, HPA052707), PMS1 (1:2000; Invitrogen, PA5-35952), PMS2 (1:1000; BD, 556415), MSH6 (1:1000; CST, 5424S), Cyclin A (1:1000; Santa Cruz, sc-56299), Tubulin (1:2500; Invitrogen, 14-4502-82) were incubated overnight at 4 °C. Secondary antibodies used were anti-rabbit HRP-linked (1:10,000-20,000; CST, 7074), anti-mouse HRP-linked (1:10,000-20,000; CST, 7076), anti-rabbit TrueBlot HRP-linked (1:1.000; Rockland,18-8816-3), anti-rabbit IRDye 800CW (1:20,000; LI-COR, 926-32213) or anti-mouse IRDye 680RD (1:20,000; LI-COR, 926-68070). Blots were imaged with Odyssey Imaging System (LI-COR) or ChemiDoc Imaging System (Bio-Rad).

#### Structural modeling

The structural modeling of MLH1 and MBD4 complex was performed using AlphaFold2 (29) with a fragmentation strategy shown to increase the sensitivity of detection (30). Sequences of MBD4 (O95243) and MLH1 (P40692) were retrieved from the UniProt database and were submitted to three iterations of MMseqs2 (31) against the uniref30_2202 database. The resulting alignments were filtered using hhfilter (32) using the parameters (‘id’=100, ‘qid’=25, ‘cov’=50) and the taxonomy assigned to every sequence, keeping only one sequence per species. Full-length sequences in alignments were then retrieved and these sequences were realigned using MAFFT (33) with the default FFT-NS-2 protocol. MLH1 and MBD4 were fragmented into 13 and 30 overlapping fragments, respectively, each of approximately 30 amino acids long in the disordered regions. To build the mixed co-alignments, sequences in the alignment of individual partners were paired according to their assigned species and left unpaired in case no common species were found. The concatenated multiple sequence alignments of tested partners were used as input of 1 run of the AlphaFold2 algorithm with 3 recycles each and 5 generated models using ColabFold v1.5.2 (34) with the Multimer v2.3 model parameters (35). Five scores were used to rate the quality of the models, the pLDDT, the pTMscore, the ipTMscore and the model confidence score (weighted combination of pTM and ipTM scores with a 20:80 ratio) together with the actifpTM score (36). 3D structures were represented using ChimeraX (37) and multiple sequence alignments were displayed using Jalview (38).

## Results

### A Cas9-based system to track the repair of cytosine deamination

To identify which DNA repair factors participate in resolving 5mC deamination in human cells, we developed an experimental framework using cytosine base editors (CBEs). CBEs are engineered CRISPR-based tools that enable precise C>T substitutions in genomic DNA without generating double-strand breaks. They typically consist of a catalytically impaired Cas9 variant fused to a cytidine deaminase enzyme, which is guided to a specific locus by a single-guide RNA (sgRNAs). Upon binding, the deaminase converts a target 5mC to T, or a target unmethylated C to U, within an exposed single-stranded editing window (25,39).

We redesigned the CBE variant YE1-BE3-FNLS, which shows high on-target editing efficiency and extremely low off-target and bystander edits (24), to encode a catalytically dead Cas9 (dCas) fused to an engineered APOBEC1. Our tool enables strand-specific cytosine deamination to produce T:G or U:G mismatches independently of DNA replication and without flanking DNA nicks, thereby mimicking spontaneous cytosine deamination in physiological conditions (Fig. 1A). Cytosine deamination within methylated CpGs in HAP1 cells confirmed C>T base-editing specifically of the target cytosine, with no detectable base-editing of the cytosine on the complementary strand (Fig. 1B).

**Figure 1.**
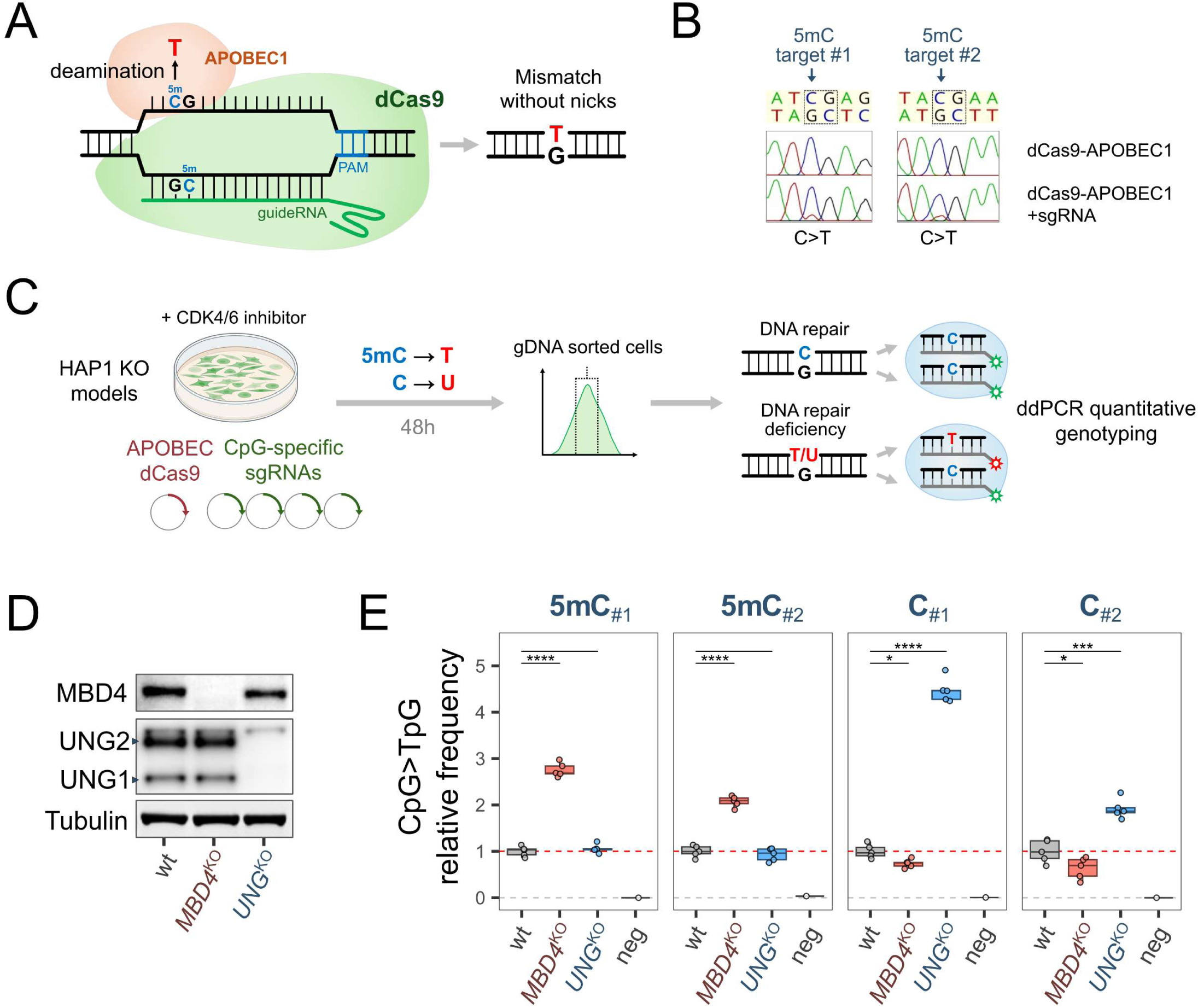
Specialized BER glycosylases repair cytosine deamination induced by a Cas9-based system. **(A)** Scheme of the cytosine base editor (CBE) used in this study, based on a catalytically dead Cas9 (dCas) fused to an engineered rat APOBEC1 cytidine deaminase. The tool induces strand-specific cytosine deamination to produce T:G or U:G mismatches within an exposed single-stranded editing window of the target sequence. **(B)** Sanger sequencing on amplified genomic DNA of HAP1 cells exposed to the CBE, with or without single-guide RNAs (sgRNA) targeting two independent methylated cytosines (5mC) loci within CpGs. **(C)** Scheme of the experimental framework used to quantify efficiency of repair following cytosine deamination, based on transfection of HAP1 cells exposed to CDK4/6 inhibitor Palbociclib with CBE and sgRNA vectors, sorting of cells of a narrow range of EGFP (co-expressed in sgRNA vectors) and mCherry (co-expressed in CBE vector), and Droplet Digital PCR (ddPCR) genotyping on genomic DNA to differentiate DNA molecules containing a T/U or C in each targeted cytosine position. **(D)** Western blotting on total cell extracts of HAP1 cells wild-type or knockout for BER glycosylases *MBD4* or *UNG.* Arrows indicate specific bands of UNG1 (mitochondrial) and UNG2 (nuclear). **(E)** ddPCR-based quantification of C>T frequency of two target 5mC loci and two unmethylated C loci within CpGs, following targeted deamination for 48h in HAP1 cells. Values are relative to the mean of wild-type replicates (shown as a dashed line). Higher C>T frequencies than wild-type cells indicate disruption of DNA repair. Five biological replicates per genotype from a single experiment are shown. Negative control (neg) represents measurements on unsorted HAP1 wild-type cells not exposed to the CBE. Two-sided unpaired *t-*test P-values without multiple comparisons adjustment are indicated following the notation: * *P* ≤ 0.05, ** *P* ≤ 0.01, *** *P* ≤ 0.001, **** *P≤* 0.0001. Non-significant P-values are not shown. Whiskers extend to the largest or lowest value up to 1.5 times the inter-quartile range from the hinges.

### dCas9-APOBEC1 validates specialized roles of BER glycosylases

Isogenic HAP1 cells harboring loss of different DNA repair components were then transiently co-transfected with vectors for dCas9-APOBEC1 and sgRNAs targeting two methylated and two unmethylated CpGs, as confirmed by bisulfite sequencing (Supplementary Fig. 1). Transfection was performed in cells exposed to Palbociclib, a selective inhibitor of the cyclin-dependent kinases CDK4/6 (40), thereby blocking cells in G_1_ and limiting fixation of mismatches as C>T transitions during S-phase. To quantify the accumulation of deaminated cytosines, we used ddPCR genotyping on genomic DNA to distinguish DNA molecules containing a T/U or C at each targeted cytosine. Notably, U has the base-pairing properties of T and is recognized as such following the end-point ddPCR reaction (Fig. 1C). In our experimental workflow, higher C>T frequencies relative to wild-type cells indicate ineffective repair of these DNA lesions.

To validate the performance of this system, we generated HAP1 cell lines knocked out for *MBD4* or *UNG*, the main BER glycosylases responsible for the repair of deaminated 5mC or unmethylated C, respectively (8,41,42) (Fig. 1D). *MBD4* knockout (^KO^) led to a 2.1-2.8-fold increase in C>T frequency at 5mC loci (Fig. 1E), consistent with previous reports relying on the endogenous rates of 5mC deamination in mammalian cells (8,43). Surprisingly, *MBD4*^KO^ led to a small but significant decrease in the C>T frequency at unmethylated C loci. MBD4 also recognizes U:G mismatches as substrates, and its loss may increase mismatch accessibility for higher-efficiency repair by UNG or SMUG1 (44). *UNG*^KO^ led to a 1.9-4.5-fold increase in C>T frequency specifically at unmethylated C loci, with no effect on the repair of deaminated 5mC loci (Fig. 1E). Of note, UNG excision efficiency depends on the DNA sequence flanking the mismatch (45), possibly explaining the variable effect of *UNG*^KO^ among unmethylated C loci.

Overall, our dCas9-APOBEC1 deamination system successfully recovered the known specialized roles of BER glycosylases in cytosine deamination repair, thereby enabling interrogation of the repair of endogenous loci in their native chromatin context.

### Non-canonical MMR plays a key role in 5mC deamination repair in coordination with MBD4

To explore the interplay of BER and MMR in 5mC deamination repair, we further disrupted different MMR complexes through knockouts of MMR members in HAP1 cells wild-type or knockout for *MBD4*. We targeted the MutLα complex (*MLH1*^KO^ or *PMS2*^KO^), MutLβ (*MLH1*^KO^ or *PMS1*^KO^) or MutSα (*MSH6*^KO^) (Fig. 2A and B). We then applied our dCas9-APOBEC1 deamination system to assess whether these DNA repair deficiencies would disrupt repair, as indicated by increased C>T frequencies relative to wild-type cells.

**Figure 2.**
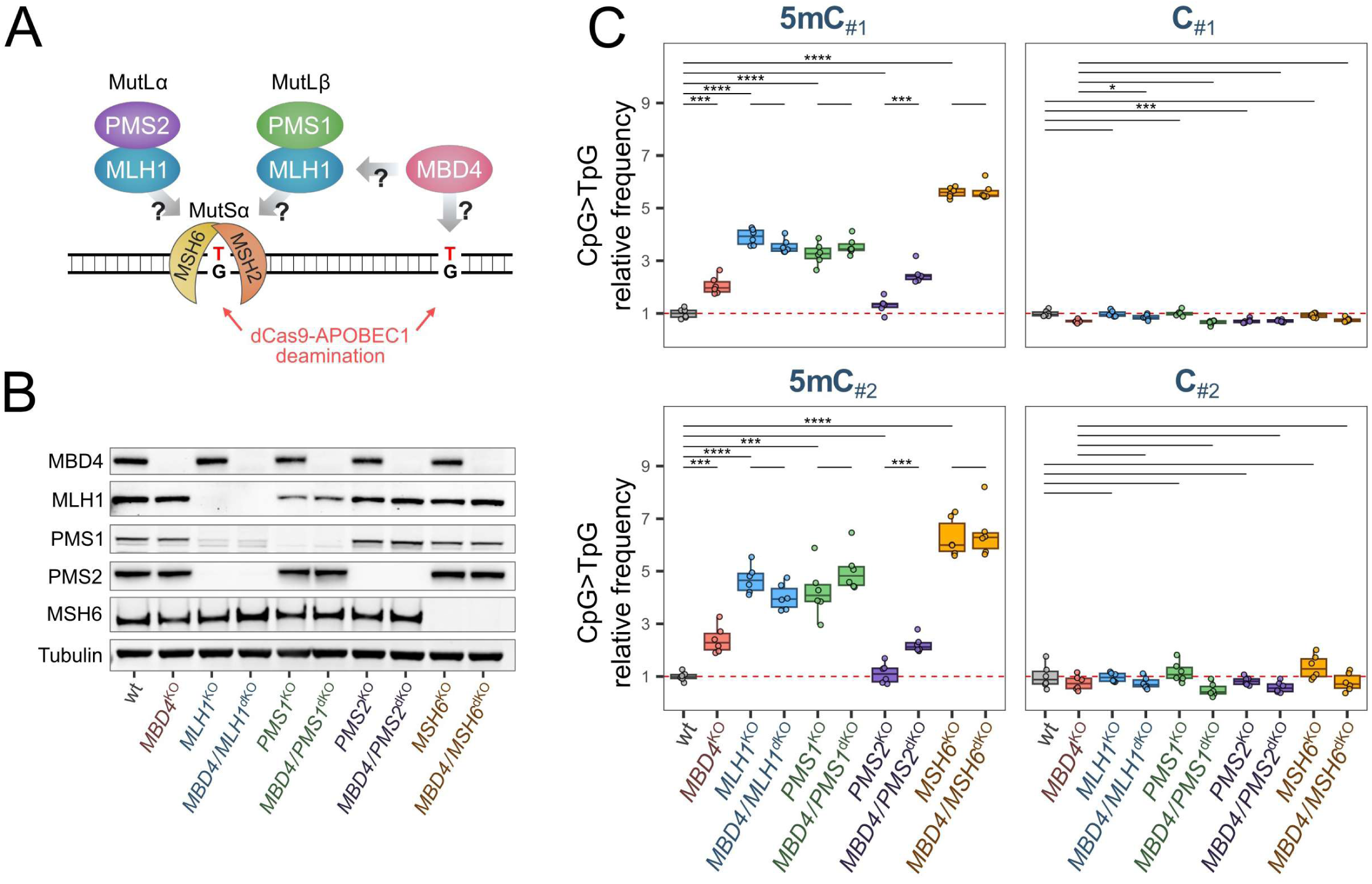
Loss of MMR members MLH1, PMS1 and MSH6 impair SmC deamination repair. **(A)** Overview of recognition of G:T mismatches caused by SmC deamination, highlighting the potential cooperative repair roles of BER glycosylase MBD4 and MMR members, including MLH1, PMS1, PMS2, MSH2 and MSH6. **(B)** Western blotting on total cell extracts of HAP1 cells wild-type or knockout for MMR members, in combination of not with *MBD4* knockout. *MLH1* knockout disrupts the stability of PMS1 and PMS2 proteins. *PMS1* knockout partially disrupts the stability of MLH1. (C) ddPCR-based quantification of C>T frequency of two target SmC loci and two unmethylated C loci within CpGs, following targeted deamination for 48h in HAP1 cells. Values are relative to the mean of wild-type replicates (shown as a dashed line). Higher C>T frequencies than wild-type cells indicate disruption of DNA repair. Three biological replicates per genotype per experiment from two independent experiments are shown. Two-sided unpaired t-test *P-*values without multiple comparisons adjustment are indicated following the notation: * *P≤* 0.05, ** *P≤* 0.01, *** *P* ≤ 0.001, **** *P≤* 0.0001. Non-significant P-values are not shown. Boxes indicate the median, 25th and 75th percentiles. Whiskers extend to the largest or lowest value up to 1.5 times the inter-quartile range from the hinges.

First, *MSH6*^KO^ led to a strong phenotype, with an increase in C>T frequency of 5.6-6.3-fold at 5mC loci (Fig. 2C), confirming the importance of upstream mismatch recognition by the MMR complex MutSα during 5mC deamination repair. To delineate the downstream complexes involved, we analyzed the following conditions. Single *MLH1*^KO^, which also disrupts the stability of PMS1 and PMS2 proteins (Fig. 2B), led to a 3.9-4.7-fold increase in C>T frequency at 5mC loci relative to wild-type cells. Single *PMS1*^KO^ led to a comparable effect to *MLH1* loss, with a 3.3-4.2-fold increase in C>T frequency at 5mC loci. In contrast, single *PMS2*^KO^ did not induce any increase in C>T frequency in comparison to wild-type cells (Fig. 2C). These data indicate that 5mC deamination repair is mediated by MutLβ, but not by MutLα-dependent post-replicative MMR. Overall, our data support the hypothesis that the MMR complex MutSα is essential for T:G mismatch recognition and initiation of repair, most likely by recruiting MutLβ.

Next, we investigated the potential crosstalk between MMR and BER glycosylase MBD4 in the repair of deaminated 5mC. Remarkably, double-knockout of *MBD4* with either *MLH1*, *PMS1* or *MSH6* led to no additive increase in C>T frequency of 5mC loci as compared to their respective single-knockouts, indicating that MBD4 activity is dependent on the MMR complexes MutLβ and MutSα. Strikingly, loss of MBD4 only partially reproduced the phenotype observed upon disruption of MutLβ and MutSα, suggesting that MBD4-independent activities of these complexes are involved in 5mC deamination repair. None of the MMR knockouts resulted in a consistent increase in C>T frequencies at unmethylated C loci relative to cells matched for *MBD4* mutational status, indicating that MMR plays a role in deamination repair specifically within methylated CpGs (Fig. 2C).

Overall, our data reveal a novel role for non-canonical MMR involving MutLβ and MutSα in 5mC deamination repair. To our knowledge, this is the first experimental evidence directly linking MMR and BER pathways as part of a coordinated DNA repair response to potentially mutagenic damage to methylated DNA.

### *PMS1* loss phenocopies the CpG>TpG hypermutation linked to *MBD4* deficiency

To orthogonally confirm the results obtained with our dCas9-APOBEC1 deamination system, we took a genome-wide approach based on WGS of long-term cultured MMR-deficient cell lines in the absence of induced DNA damage, thus relying on endogenous deamination rates. Single-cell clonal lines were cultured for 60-90 days to permit mutation accumulation, followed by a second single-cell cloning and WGS of 2-4 individual subclones (Fig. 3A). We excluded variants present in either parental HAP1 cells or shared among subclones, thereby focusing on mutations that occurred after the isolation of the initial knockout clones. This dataset was complemented by mutational profiles we previously obtained using a similar strategy in *MBD4*^KO^ versus wild-type HAP1 cells (8). Data from wild-type HAP1 subclones were used as a reference for the background mutagenesis arising from *in vitro* culturing (46).

**Figure 3.**
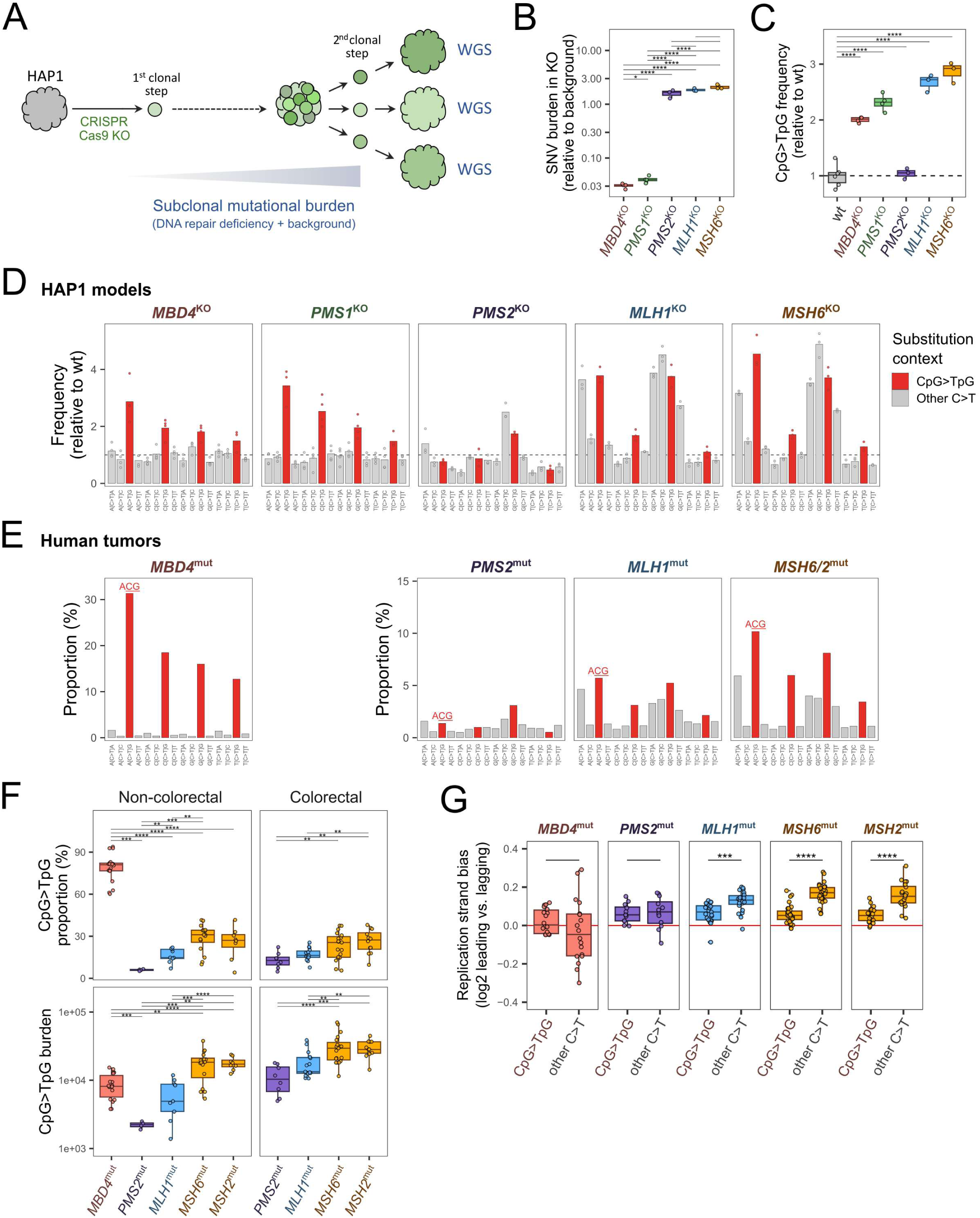
Genome-wide CpG>TpG hypermutagenesis of MMR-deficient cells and tumors is partly linked to SmC deamination. **(A)** Workflow used to quantify single nucleotide variant (SNV) mutational burden of isogenic cell lines. Expanded clones of diploidized HAP1 cells wild-type or knockout for MMR genes were long-term cultured, followed by a second subcloning step, expansion and WGS of multiple subclones per genotype. Subclone-specific mutational burdens were quantified, representing a combination of knockout-related mutations due to specific DNA repair deficiencies and background mutations due to *in vitro* culturing. **(B)** Mutational signature analysis of subclones to linearly estimate the mutational burdens specifically due to knockouts of DNA repair genes or background. The background mutational signature used for fitting was obtained from wild-type subclones. Data for all SNV classes are shown. Two-sided unpaired *t-test* P-values without multiple comparisons adjustment are indicated. **(C)** Distributions of CpG>TpG substitution frequencies among all SNVs, as obtained by WGS in HAP1 subclones, relative to the mean of wild-type subclones (shown as a dashed line). Two-sided unpaired *t-test* P-values without multiple comparisons adjustment are indicated. **(D)** C > T substitution frequencies by trinucleotide context among all SNVs, relative to the mean of wild-type subclones (shown as a dashed line). Bars represent the means of subclones. **(E)** C > T substitution percentages by trinucleotide context among all SNVs of human tumor samples of non-colorectal origin. Bars represent the means among cases harboring germline or somatic biallelic mutations in *MBD4* (n=18), *PMS2* (n=4), *MLH1* (n=9) or *MSH2/6* (n=24). **(F)** Distributions of CpG>TpG percentages among all SNVs or absolute CpG>TpG burdens of human tumor samples of either colorectal or non-colorectal origins. Two-sided Wilcoxon test P-values are indicated. Of all possible pairwise comparisons, only significant ones are shown. **(G)** Replication strand asymmetry bias in the leading versus the lagging strands among C>T mutations. Tumor samples of both colorectal and non-colorectal origins were grouped. No bias for either strand is indicated as a red line. Two-sided Wilcoxon test P-values are indicated, and non-significant P-values are not shown. Statistics shown throughout the figure follow the notation: * *P* ≤ 0.05, ** *P* ≤ 0.01, *** *P* ≤ 0.001, **** *P* ≤ 0.0001. Boxes indicate the median, 25th and 75th percentiles. Whiskers extend to the largest or lowest value up to 1.5 times the inter-quartile range from the hinges.

The consequences of post-replicative repair dysfunctions for small insertions and deletions (InDels) have been previously described in detail (47). We therefore focused our analysis on SNVs. We applied mutational signature analysis on SNVs to linearly estimate mutational burden linked to each gene knockout (Supplementary Fig. 2). *PMS2*^KO^, *MLH1*^KO^ and *MSH6*^KO^ were associated with strong SNV hypermutated profiles, consistent with disruption of post-replicated MMR in highly proliferative HAP1 cells (Fig. 3B). In contrast, *MBD4*^KO^ and *PMS1*^KO^ led to a more focused phenotype, characterized by a low SNV mutational burden but highly enriched in CpG>TpG mutations (Fig. 3B and C; Supplementary Fig. 3). Consistent with data obtained with our dCas9-APOBEC1 deamination system, *MBD4*^KO^ was associated with a 2-fold increase in CpG>TpG relative frequency, while *PMS1*^KO^ was associated with a significantly higher effect of 2.3-fold (two-sided unpaired *t*-test; *P*=0.021). Remarkably, CpG>TpG mutations in *PMS1*^KO^ cells showed similar relative frequencies by trinucleotide context to *MBD4*^KO^ cells (Fig. 3D), with the pattern A<u>CG</u>>C<u>CG</u>>G<u>CG</u>>T<u>CG</u> characteristic of the SBS96 mutational signature of *MBD4*-deficient human tumors (8,28). Altogether, we show that *PMS1* loss phenocopies the preferential CpG>TpG hypermutator profile of *MBD4*-deficient cells, indicating that 5mC deamination repair may represent one of the main functions of human PMS1.

The relative frequency of CpG>TpG mutations in *PMS2*^KO^ cells was comparable to wild-type cells (two-sided unpaired *t*-test; *P*=0.69), indicating that disruption of post-replicative MMR does not lead to preferential CpG mutagenesis (Fig. 3C). This pattern is fully consistent with that obtained with our dCas9-APOBEC1 deamination system (Fig. 2C). In contrast, both *MLH1*^KO^ and *MSH6*^KO^ led to a robust increase in CpG>TpG relative frequency (2.7 and 2.9-fold, respectively), although C>T transitions in other trinucleotide contexts were also highly enriched (Fig. 3C and D). Notably, *MLH1*^KO^ and *MSH6*^KO^ cells showed the highest C>T peak within the GCC context and a large contribution of T>C mutations, similarly to post-replicative MMR disruption through *PMS2*^KO^ (Fig. 3C and D; Supplementary Fig. 3). Overall, our genome-wide data support our hypothesis that while PMS2 is primarily essential for post-replicative MMR, MLH1 and MSH6 play central roles in both 5mC deamination repair and post-replicative MMR.

It is important to state that the genome-wide approach we employed cannot fully untangle CpG mutagenesis driven by failed 5mC deamination repair or processes linked to DNA replication, particularly due to the significant background mutational burden due to *in vitro* culturing. However, our data are consistent with the patterns observed with our dCas9-APOBEC1 deamination system, in which mismatches are introduced in the genome independently of DNA replication. More importantly, we orthogonally confirmed that PMS1 plays a major role in protecting the genome from CpG>TpG mutagenesis.

### 5mC deamination contributes to the CpG>TpG burden of tumors deficient for *MLH1*, *MSH6 and MSH2*

We next wondered whether the mutational patterns of MMR-deficient human tumors would support the involvement of MMR factors in 5mC deamination repair. We reanalyzed whole genomes from the Genomics England pan-cancer series (GEL series release data v17; 12,726 high-quality tumor samples) (50), in addition to a collection of two in-house and four previously described *MBD4*-deficient tumors (43,51). Tumors deficient for *MBD4* in the GEL series have been previously identified and extensively characterized (8). We additionally identified MMR-deficient tumors by performing mutational signature analysis (28), excluding samples with overlapping DNA polymerase defects, and focusing on those harboring germline or biallelic somatic loss-of-function mutations in *PMS2*, *MLH1*, *MSH6* or *MSH2*. *PMS1* loss in human cancer remains anecdotal and previously described cases lack sequencing data on tumor samples (48,49).

The relative frequencies of CpG>TpG mutations across trinucleotide contexts in non-colorectal human tumors closely mirrored those observed in HAP1 cells. Preferential C>T mutagenesis at the A<u>CG</u> context, indicative of 5mC deamination repair deficiency (e.g. *MBD4* and *PMS1* loss), was minimally present in *PMS2*-inactivated tumors, but was increasingly prominent in *MLH1*-inactivated and *MSH6/MSH2*-inactivated tumors, respectively (Fig. 3E; Supplementary Fig. 4). Furthermore, among MMR-deficient tumors of non-colorectal origin, we observed the lowest CpG>TpG percentages and absolute burden in *PMS2*-inactivated tumors (6.0 ± 0.6%; 2219 ± 255), followed by *MLH1*-inactivated tumors (16.2 ± 5.0%; 6229 ± 3643), and *MSH6/MSH2*-inactivated tumors (27.7 ± 10.3%; 17816 ± 7380). The same overall pattern was observed for tumors of colorectal origin, which are associated with higher burden of all SNVs, including CpG>TpG transitions (Fig. 3F; Supplementary Fig. 5).

Previous studies have shown that defects in the MMR pathway cause asymmetric mismatch patterns arising during leading- and lagging-strand DNA synthesis (52). In contrast, mutagenesis driven by spontaneous 5mC deamination does not exhibit replication-strand bias, as observed in *MBD4*-deficient tumors (Fig. 3G). Strikingly, tumors lacking *MLH1*, *MSH6* or *MSH2* exhibited a weaker replication-strand bias toward the leading strand for CpG>TpG mutations compared with other C>T transitions (Fig. 3G). This pattern suggests that the elevated CpG>TpG load in these tumors is at least partly due to failed repair of 5mC deamination rather than defects in post-replicative MMR. In contrast, *PMS2*-mutated tumors showed similar levels of replication-strand bias for CpG>TpG and other C>T transitions (Fig. 3G). Overall, the mutational landscape of MMR-deficient tumors aligns with our experimental results, demonstrating that MutLβ and MutSα, but not MutLα, play major roles in 5mC deamination repair.

### MBD4 engages with the MutLβ complex by directly binding to MLH1

MBD4 has been shown to form a stable complex with MLH1 and PMS2, although experiments using a co-expression baculovirus system indicated a direct physical interaction only between MBD4 and MLH1, and not between MBD4 and PMS2 (18,19,53,54). Given that our data do not support a role for PMS2 in the repair of 5mC deamination, we next considered whether MBD4 might assemble into a multimeric complex with MLH1 and PMS1 to coordinate the repair response to this type of DNA damage.

To test this hypothesis, we generated HAP1 cell lines stably overexpressing MBD4 constructs bearing either C- or N-terminal FLAG tags and performed co-immunoprecipitation assays. We successfully co-immunoprecipitated MLH1, PMS1 and PMS2 with both MBD4 constructs as baits, while the interactions were consistently stronger with the N-terminally tagged MBD4. Comparable interaction levels were detected in asynchronous and G_1_-arrested cells, indicating that MBD4 association with these MMR components is largely independent of cell cycle phase regulation (Fig. 4A). We further confirmed the interaction between MBD4 and the MutLβ complex by immunoprecipitating endogenous MBD4 from wild-type untagged HAP1 cell extracts using PMS1- and MLH1-specific antibodies (Fig. 4B).

**Figure 4.**
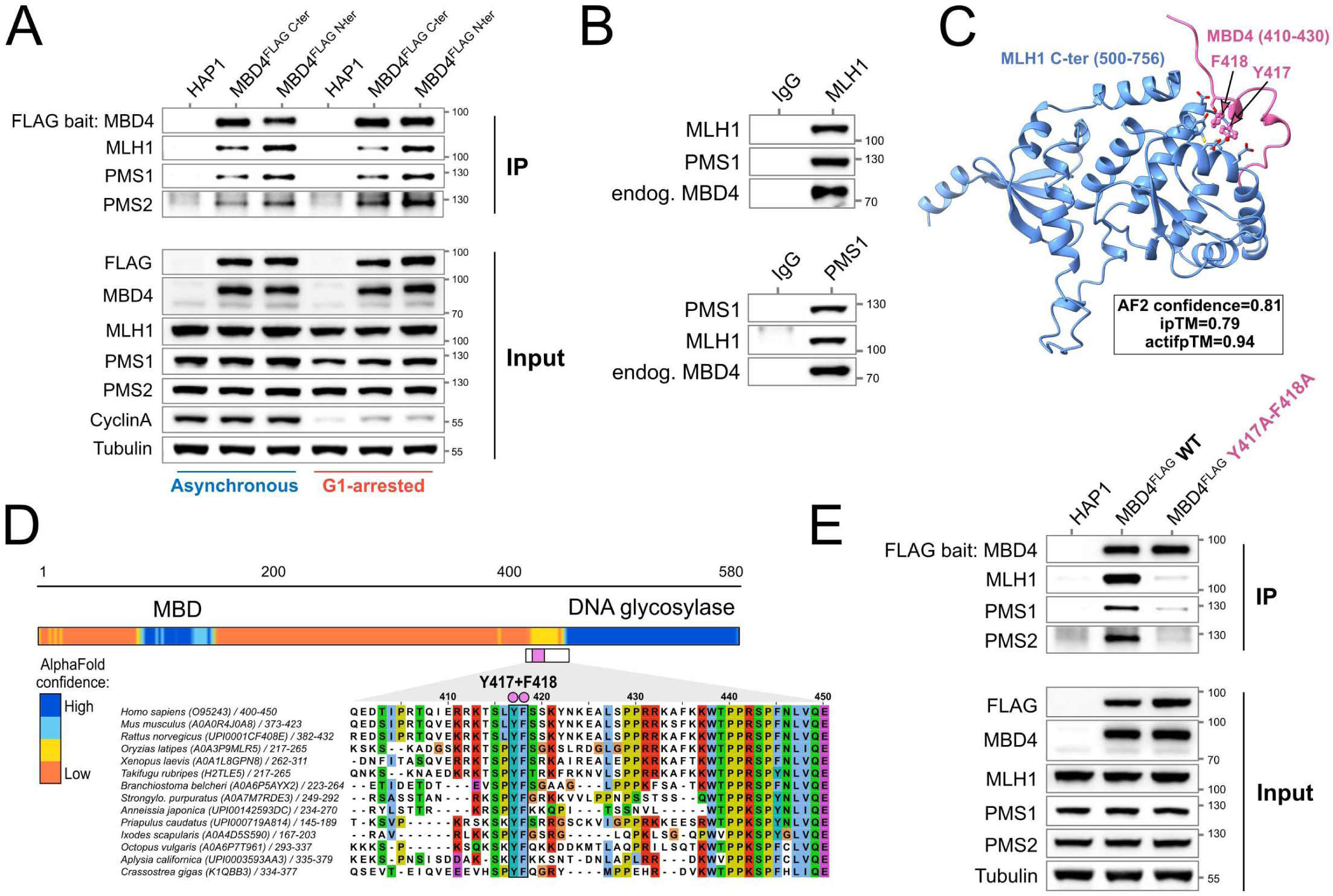
MBD4 physically interacts with the Mutll3 components MLH1 and PMS1. **(A)** Western blotting of anti-FLAG immunoprecipitated HAP1 total cell extracts, and respective inputs. Parental cells or stably overexpressing C- or N-terminally FLAG-tagged MBD4 and were cultured with vehicle (asynchronous) or with 1 µM Palbociclib (G1-arrested) for 17h. Cyclin A was used as a marker of cycling cells. **(B)** Western blotting of immunoprecipitated total cell extracts of non-tagged parental HAP1 cells, using antibodies specific for MLH1, PMS1 or isotype lgG control for immunoprecipitation. **(C)** AlphaFold2 model of the MLH1 C-terminal domain in complex with MBD4. Ribbon representation of the MLH1 C-terminal domain (residues 500-756) shown in blue, in complex with a C-terminal fragment of MBD4 (residues 410-430) shown in pink. The interaction interface is highlighted, with MBD4 residues Y417 and F418 displayed as sticks and indicated by arrows. These conserved aromatic residues form a key contact surface with MLH1. The model was generated using AlphaFold2, with an overall confidence score of 0.81, an ipTM score of 0.79, and an actual interface pTM (actifpTM) score of 0.94, indicating a high-confidence interface prediction. **(D)** Evolutionary conservation of the MLH1-interacting region in MBD4. Schematic representation of human MBD4 highlighting the methyl-CpG binding domain (MBD) and the DNA glycosylase domain, together with AlphaFold confidence coloring along the protein sequence (high confidence in blue, low confidence in orange). A multiple sequence alignment of the MBD4 C-terminal region (around residues 400-450) from representative metazoan species is shown below. The conserved aromatic residues Y417 and F418 (in human) which mediate interaction with the MLH1 C-terminal domain, are highlighted. This region shows strong conservation across vertebrates and invertebrates, supporting an evolutionarily conserved MLH1-MBD4 interaction interface. **(E)** Western blotting of anti-FLAG immunoprecipitated HAP1 total cell extracts, and respective inputs. Parental cells or stably overexpressing N-terminally FLAG-tagged MBD4, either wild-type or harboring Y417A and F418A point mutations, were used.

Next, we used a scanning protocol based on AlphaFold2 Multimer (29,30) to generate a structural model of the direct interaction between MBD4 and MLH1. AlphaFold2 predicted a high-confidence interaction between the C-terminal region of MLH1 and a highly conserved motif in MBD4 encompassing residues Y417 and F418 (confidence score of 0.81) (Fig. 4C and D, Supplementary Fig. 6). This short motif lies immediately adjacent to the DNA glycosylase domain but is predicted to be intrinsically disordered in the absence of MLH1 (Fig. 4D).

Based on our predicted 3D interaction map, we examined the contact interface between MBD4 and MLH1 by introducing point mutations into residues predicted to contribute to complex formation. To this end, we generated HAP1 cell lines stably expressing N-terminally FLAG-tagged MBD4, either wild-type or carrying Y417A/F418A variants. Co-IP assays showed that these mutations resulted in near-complete loss of binding to MLH1, PMS1 and PMS2 (Fig. 4E). Consistent with prior reports that MLH1 functions as the central scaffold for MutLα and MutLβ complex assembly, our data further suggest that the interaction between MBD4 and PMS1 is largely MLH1-dependent. Most importantly, our data shed light on the possible structural basis for the role of MutLβ in 5mC deamination repair in coordination with MBD4.

## Discussion

Spontaneous deamination of 5mC is a major endogenous source of CpG>TpG mutations involved vertebrate genomic evolution, cancer and aging. In this study, by redesigning a high-fidelity cytosine base editor to generate T:G or U:G mismatches independently of DNA replication, we created a system that closely mimics spontaneous cytosine deamination in native chromatin. We validated the sensitivity and specificity of this dCas9-APOBEC1 deamination system and confirmed that it can disentangle parallel repair activities by specialized BER glycosylases at individual cytosines in bulk cell populations. We extend on these findings by showing that T:G mismatches arising from 5mC deamination are processed by a pathway that couples the BER glycosylase MBD4 to non-canonical MMR involving MutSα (MSH2-MSH6) and MutLβ (MLH1-PMS1), rather than MutLα (MLH1-PMS2)-dependent post-replicative MMR.

We show that an intrinsically disordered domain in MBD4 is essential for its physical interaction with MutLβ, thus providing a structural basis for the convergence of BER and MMR in 5mC deamination repair. Notably, this region was omitted from earlier crystal structures of the MBD4 catalytic domain, which resolved only the folded glycosylase core (55,56). Our working model, supported by recent biochemical evidence (19), suggests that MLH1 binding to this motif may activate or enhance MBD4 glycosylase activity.

The ability of post-replicative MMR to target repair to the newly synthesized DNA strand is crucial to preserve the genetic information to the next cell generation. During DNA replication, MMR corrects mismatches errors in a strand-specific manner, with strand discrimination directed either by pre-existing strand breaks (such as the ends of Okazaki fragments) or by PCNA-guided endonucleolytic cleavage mediated by MutLα (57). In contrast, non-canonical MMR acting in clustered DNA lesions refractory to repair by DNA glycosylases are believed to be largely independent of DNA replication, lack strand directionality, and promote the recruitment of the error-prone polymerase-η to chromatin (16,17,58). However, at isolated deaminated 5mC sites, MBD4 may guide MMR toward the damaged DNA strand to promote error-free repair. This model is consistent with MBD4 highly specialized function in directing specific thymine excision following 5mC deamination (55).

Although recent evidence suggests a role of PMS1 in repeat expansion (59-61), the MutLβ heterodimer is dispensable for post-replicative MMR (62). We show that *PMS1* loss phenocopies the CpG>TpG hypermutated phenotypes associated with *MBD4* deficiency, suggesting that 5mC deamination repair is a major function of human PMS1. While recurrent homozygous and heterozygous *MBD4* deficiency are linked to cancer predisposition syndromes (21,22), very little evidence supports a role of *PMS1* in cancers (48,49), for unclear reasons that need to be addressed in the future.

In summary, our study delineates a previously unrecognized MBD4-MutLβ-driven repair pathway that preserves the integrity of methylated cytosines, establishing non-canonical MMR as a fundamental safeguard of the human genome against 5mC deamination-induced mutagenesis.

---

**Supplementary Figure 1.**
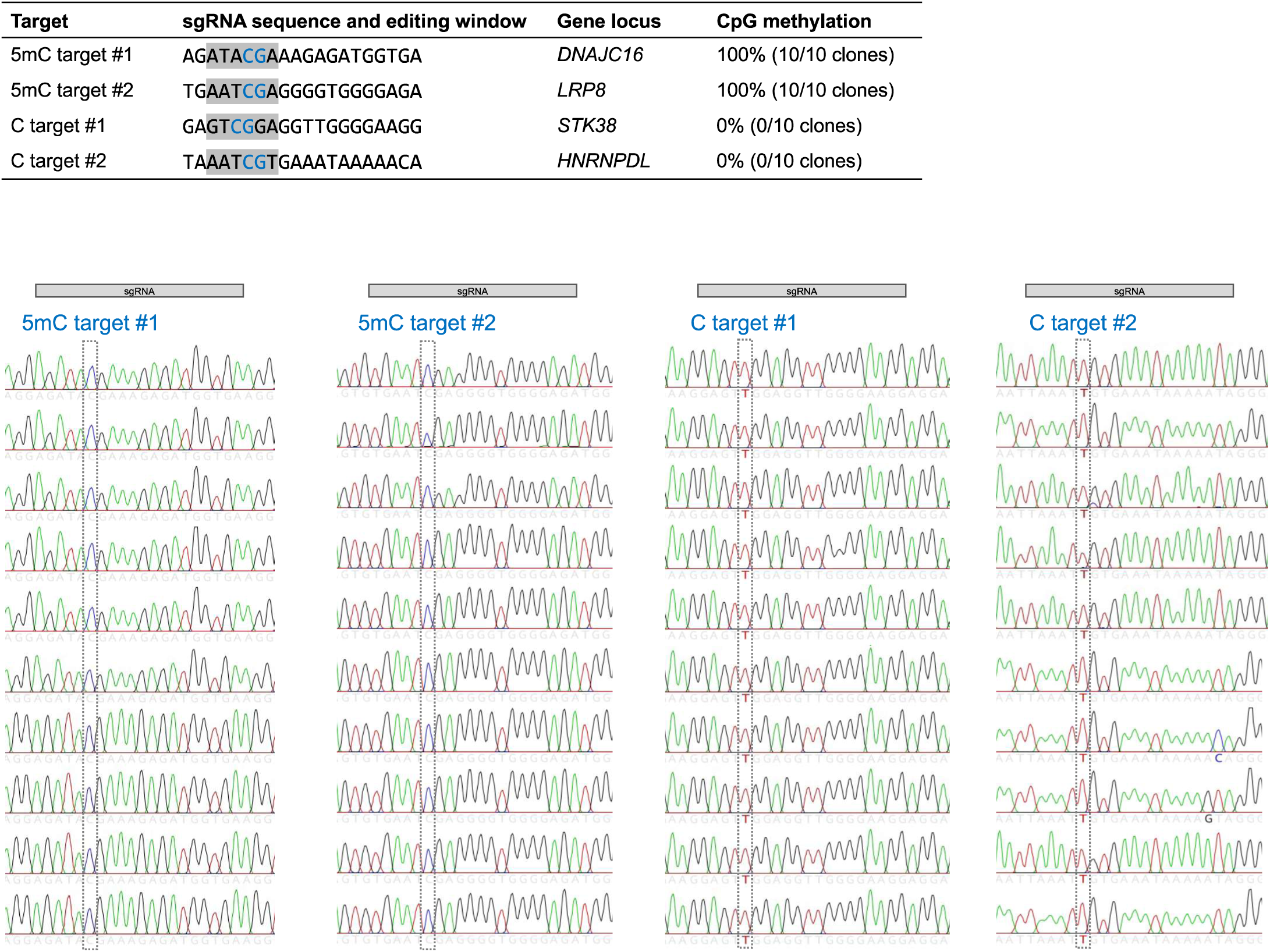
Methylation status of cytosine loci targeted with the Cas9 deamination system. Single-guide RNA target sequences are indicated, including the dCas9-APOBEC1 editing window in grey, and target CpG in blue (above). Sanger sequencing on cloned fragments of amplified bisulfite-converted genomic DNA of wild-type HAP1 cells. Sequencing of 10 individual clones per target sequence are shown. Bisulfite converts unmethylated cytosines to thymines, as shown for unmethylated C targets #1 and #2 (below).

**Supplementary Figure 2.**
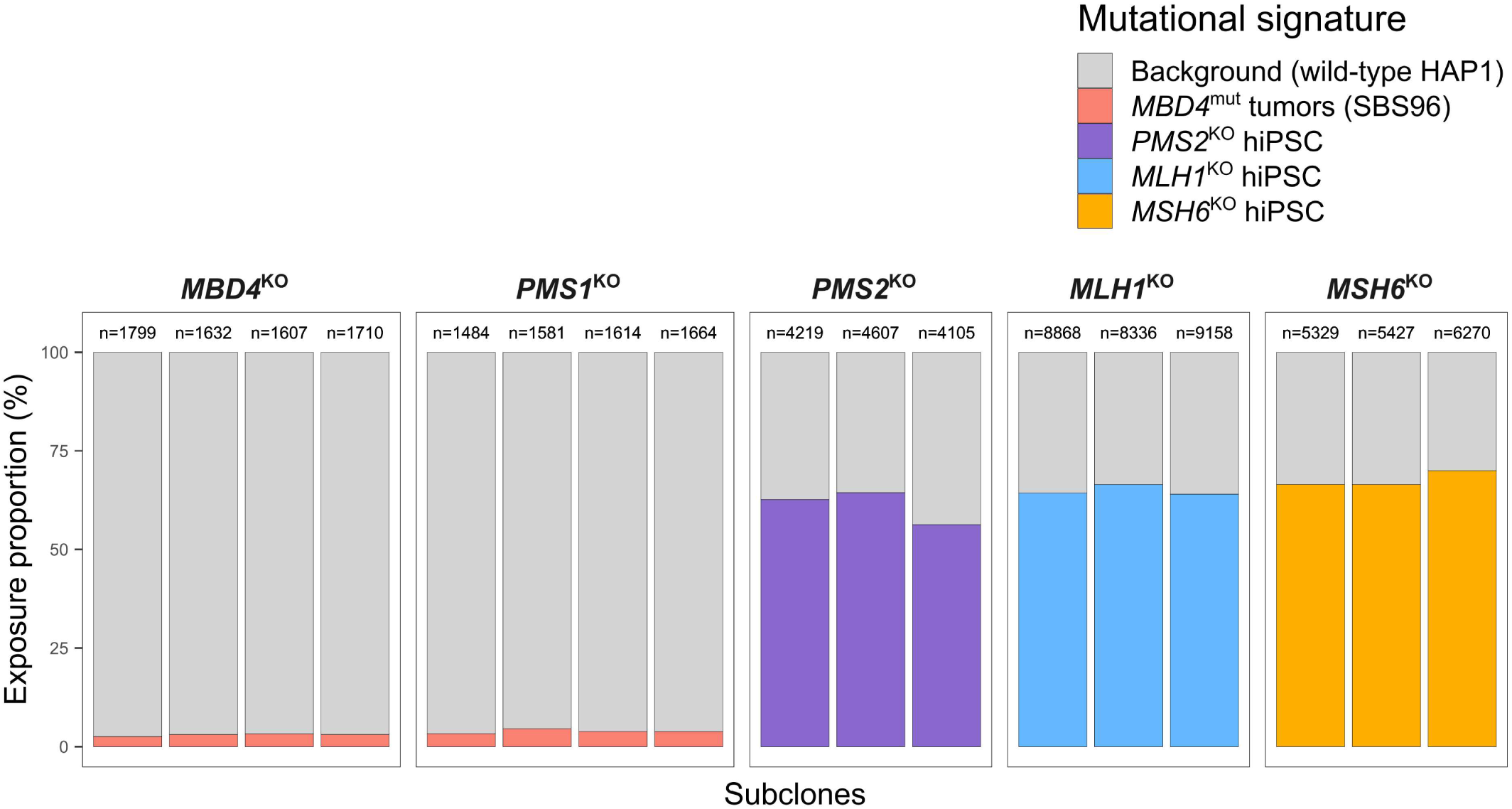
Mutational signature analysis of HAP1 subclones analyzed by WGS. Exposure to MMR-deficiency signatures previously obtained in human Induced pluripotent stem cell (hiPSC), single base substitution signature SBS96 of MBĐ4-deficient human tumors, or background mutational signature derived from the mean mutational profiles of wild-type HAP1 subclones. Total number of SNVs per subclone are indicated.

**Supplementary Figure 3.**
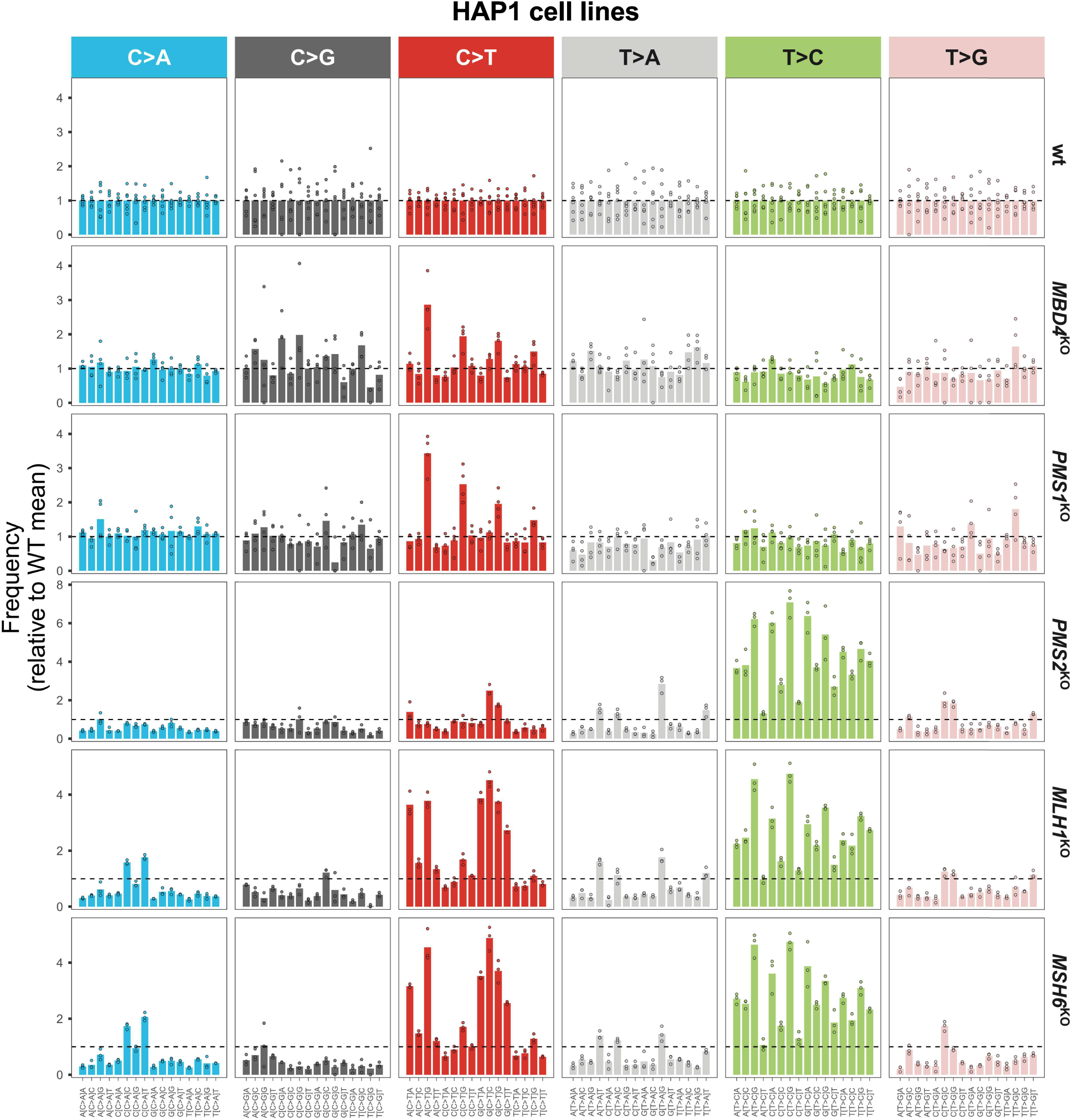
Substitution profiles by trinucleotide context of *MBD4* and MMR genes knockout HAP1 cell models. Distributions of substitution frequencies obtained by WGS in HAP1 subclones wild-type or knock­out for *MBD4, PMS1, PMS2, MLH1* and *MSH6.* Values shown are relative to the mean of wild-type subclones, shown as a dashed line. Bars represent the mean of subclones, shown as individual points.

**Supplementary Figure 4.**
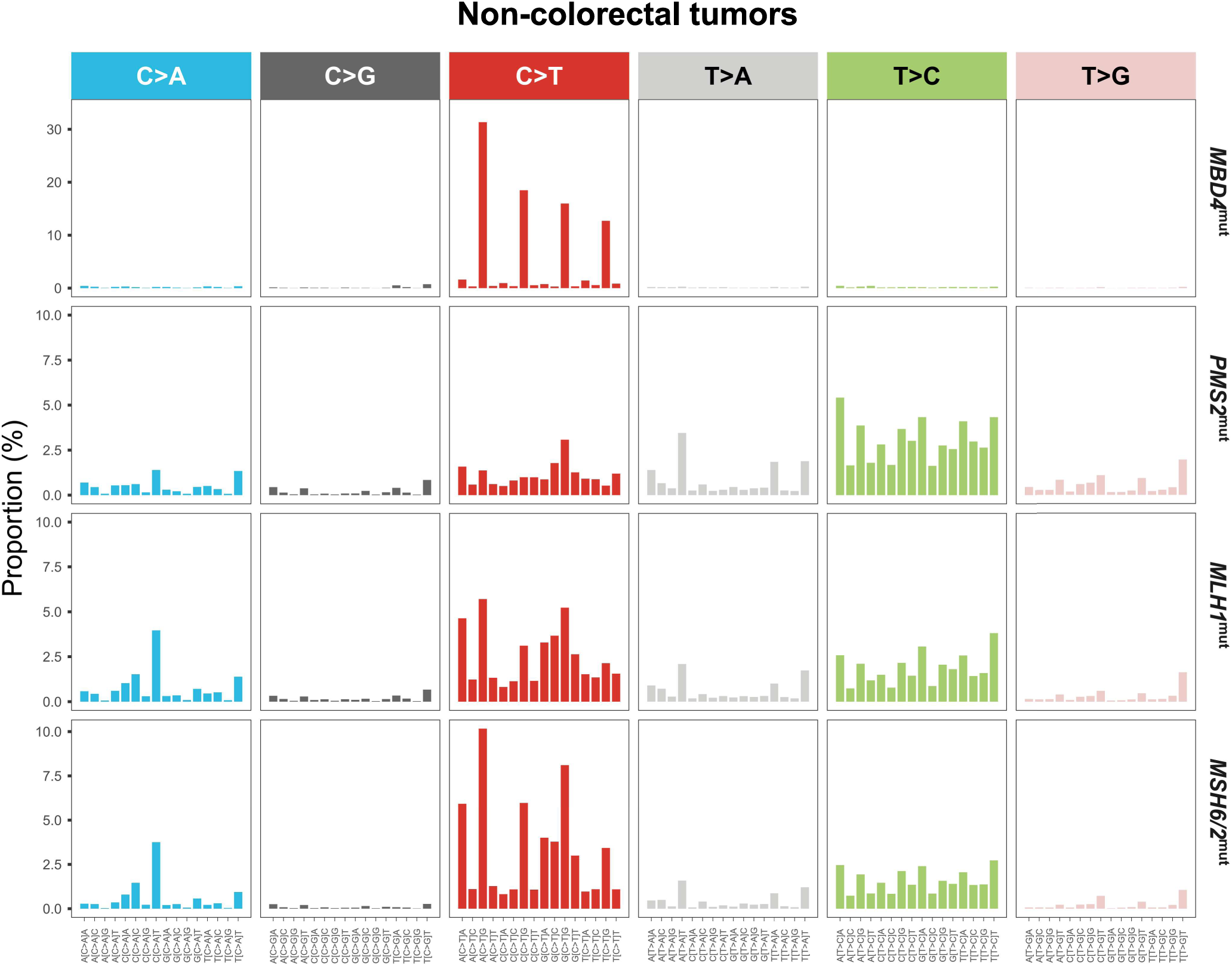
Substitution profiles by trinucleotide context of MBD4 and MMR genes-mutated human tumors. Bars represent the means among tumors of non-colorectal origin, harboring germline or somatic biallelic mutations in *MBD4* (n=18), *PMS2* (n=4), *MLH1* (n=9) or *MSH2/6* (n=24).

**Supplementary Figure 5.**
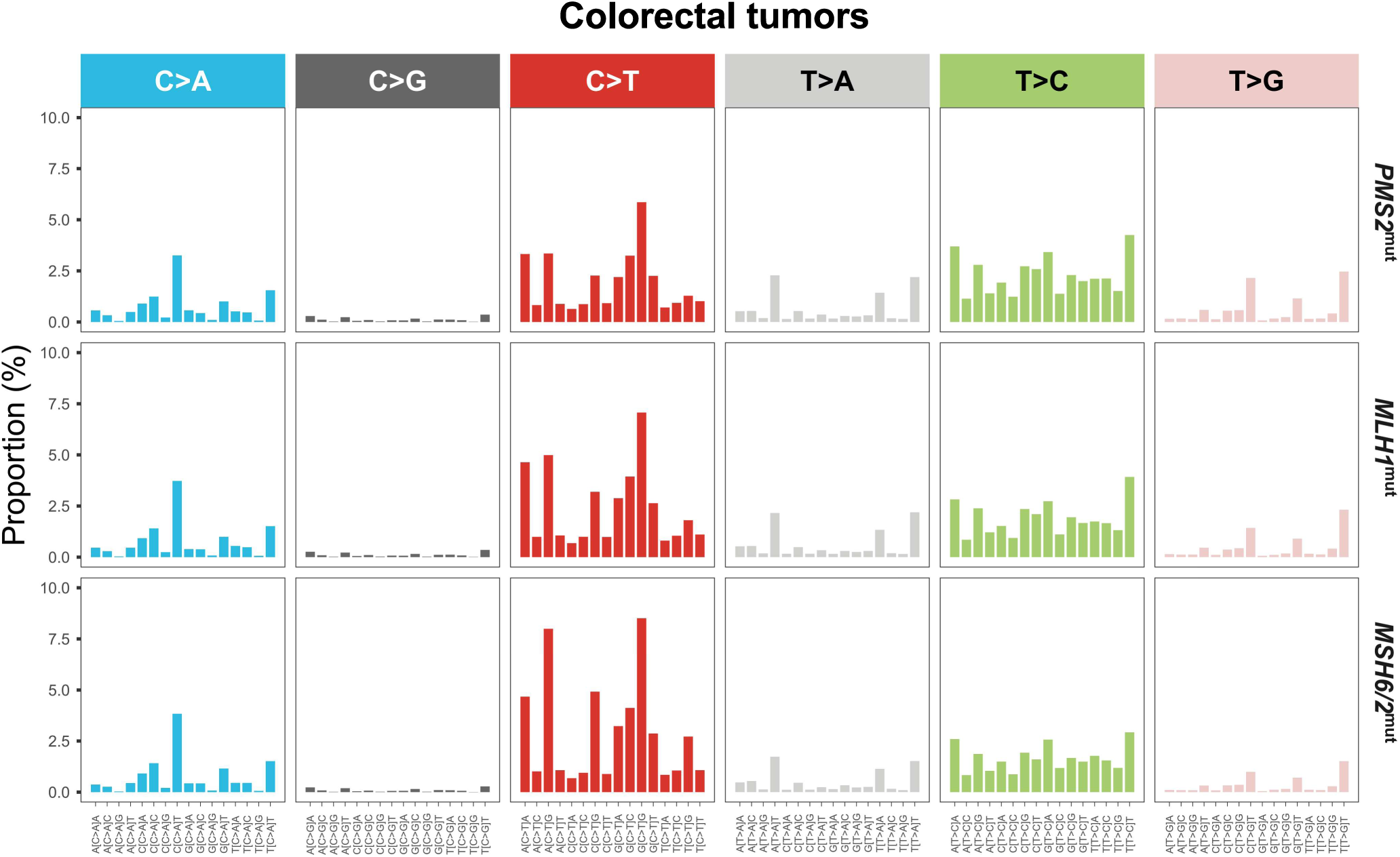
Substitution profiles by trinucleotide context of MBD4 and MMR genes-mutated human tumors. Bars represent the means among tumors of colorectal origin, harboring germline or somatic biallelic mutations in *PMS2* (n=8), *MLH1* (n=15) or *MSH2/6* (n=31).

**Supplementary Figure 6.**
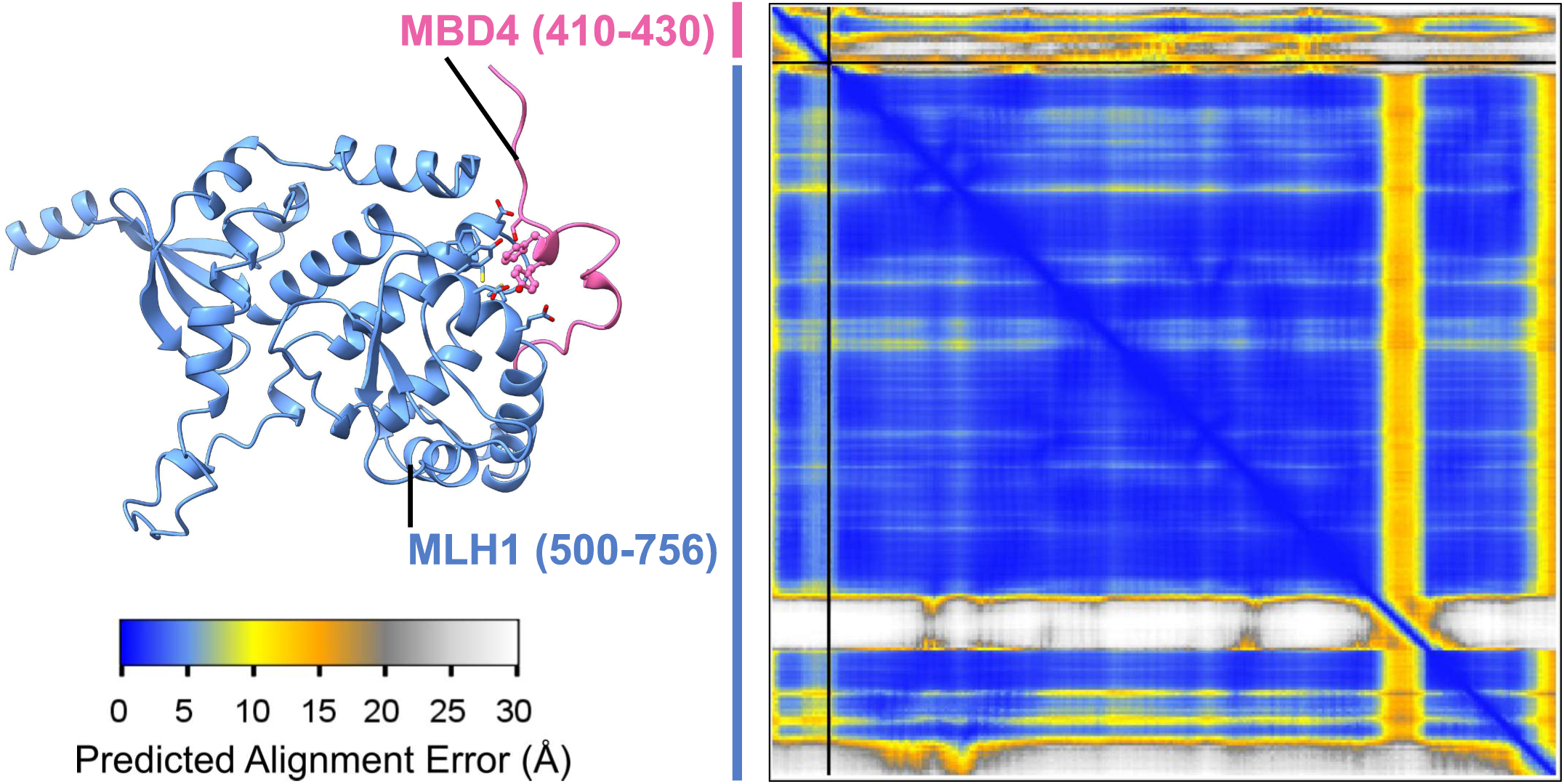
Confidence of the MBD4-MLH1 interaction through the Predicted Alignment Error map. Predicted Alignment Error (PAE) matrix colored according to the predicted alignment error scale (in Å) indicating high confidence in the relative positioning of the two proteins within the complex. Blue regions indicate low predicted error and high confidence in the relative orientation of residues whereas yellow to white regions indicate higher uncertainty. The low PAE values at the interface between MBD4 (residues 410-430, pink) and the C-terminal region of and MLH1 (residues 500-756, blue) support a well-defined and confident interaction between these regions.

## Notes

### Competing Interest Statement

The authors have declared no competing interest.

## References

1. Suzuki, M.M. and Bird, A. (2008) DNA methylation landscapes: provocative insights from epigenomics. Nat Rev Genet, 9, 465–476.

2. Hendrich, B., Hardeland, U., Ng, H.H., Jiricny, J. and Bird, A. (1999) The thymine glycosylase MBD4 can bind to the product of deamination at methylated CpG sites. Nature, 401, 301–304.

3. Blokzijl, F., de Ligt, J., Jager, M., Sasselli, V., Roerink, S., Sasaki, N., Huch, M., Boymans, S., Kuijk, E., Prins, P. et al. (2016) Tissue-specific mutation accumulation in human adult stem cells during life. Nature, 538, 260–264.

4. Kong, A., Frigge, M.L., Masson, G., Besenbacher, S., Sulem, P., Magnusson, G., Gudjonsson, S.A., Sigurdsson, A., Jonasdottir, A., Jonasdottir, A. et al. (2012) Rate of de novo mutations and the importance of father’s age to disease risk. Nature, 488, 471–475.

5. Alexandrov, L.B., Jones, P.H., Wedge, D.C., Sale, J.E., Campbell, P.J., Nik-Zainal, S. and Stratton, M.R. (2015) Clock-like mutational processes in human somatic cells. Nat Genet, 47, 1402–1407.

6. Alexandrov, L.B., Nik-Zainal, S., Wedge, D.C., Aparicio, S.A., Behjati, S., Biankin, A.V., Bignell, G.R., Bolli, N., Borg, A., Borresen-Dale, A.L. et al. (2013) Signatures of mutational processes in human cancer. Nature, 500, 415–421.

7. Krokan, H.E. and Bjoras, M. (2013) Base excision repair. Cold Spring Harb Perspect Biol, 5, a012583.

8. Silveira, A.B., Houy, A., Ganier, O., Ozemek, B., Vanhuele, S., Vincent-Salomon, A., Cassoux, N., Mariani, P., Pierron, G., Leyvraz, S. et al. (2024) Base-excision repair pathway shapes 5-methylcytosine deamination signatures in pan-cancer genomes. Nat Commun, 15, 9864.

9. Rodrigues, M., Mobuchon, L., Houy, A., Alsafadi, S., Baulande, S., Mariani, O., Marande, B., Ait Rais, K., Van der Kooij, M.K., Kapiteijn, E. et al. (2019) Evolutionary Routes in Metastatic Uveal Melanomas Depend on MBD4 Alterations. Clin Cancer Res, 25, 5513–5524.

10. Rodrigues, M., Mobuchon, L., Houy, A., Fievet, A., Gardrat, S., Barnhill, R.L., Popova, T., Servois, V., Rampanou, A., Mouton, A. et al. (2018) Outlier response to anti-PD1 in uveal melanoma reveals germline MBD4 mutations in hypermutated tumors. Nat Commun, 9, 1866.

11. Saint-Ghislain, M., Derrien, A.C., Geoffrois, L., Gastaud, L., Lesimple, T., Negrier, S., Penel, N., Kurtz, J.E., Le Corre, Y., Dutriaux, C. et al. (2022) MBD4 deficiency is predictive of response to immune checkpoint inhibitors in metastatic uveal melanoma patients. Eur J Cancer, 173, 105–112.

12. Johansson, P.A., Stark, A., Palmer, J.M., Bigby, K., Brooks, K., Rolfe, O., Pritchard, A.L., Whitehead, K., Warrier, S., Glasson, W. et al. (2019) Prolonged stable disease in a uveal melanoma patient with germline MBD4 nonsense mutation treated with pembrolizumab and ipilimumab. Immunogenetics, 71, 433–436.

13. Jiricny, J. (2006) The multifaceted mismatch-repair system. Nat Rev Mol Cell Biol, 7, 335–346.

14. Kadyrov, F.A., Dzantiev, L., Constantin, N. and Modrich, P. (2006) Endonucleolytic function of MutLalpha in human mismatch repair. Cell, 126, 297–308.

15. Crouse, G.F. (2016) Non-canonical actions of mismatch repair. DNA Repair (Amst), 38, 102–109.

16. Zlatanou, A., Despras, E., Braz-Petta, T., Boubakour-Azzouz, I., Pouvelle, C., Stewart, G.S., Nakajima, S., Yasui, A., Ishchenko, A.A. and Kannouche, P.L. (2011) The hMsh2-hMsh6 complex acts in concert with monoubiquitinated PCNA and Pol eta in response to oxidative DNA damage in human cells. Mol. Cell, 43, 649–662.

17. Pena-Diaz, J., Bregenhorn, S., Ghodgaonkar, M., Follonier, C., Artola-Boran, M., Castor, D., Lopes, M., Sartori, A.A. and Jiricny, J. (2012) Noncanonical mismatch repair as a source of genomic instability in human cells. Mol. Cell, 47, 669–680.

18. Bellacosa, A., Cicchillitti, L., Schepis, F., Riccio, A., Yeung, A.T., Matsumoto, Y., Golemis, E.A., Genuardi, M. and Neri, G. (1999) MED1, a novel human methyl-CpG-binding endonuclease, interacts with DNA mismatch repair protein MLH1. Proc Natl Acad Sci U S A, 96, 3969–3974.

19. Papin, C., Ibrahim, A., Sabir, J.S.M., Le Gras, S., Stoll, I., Albiheyri, R.S., Zari, A.T., Bahieldin, A., Bellacosa, A., Bronner, C. et al. (2023) MBD4 loss results in global reactivation of promoters and retroelements with low methylated CpG density. J Exp Clin Cancer Res, 42, 301.

20. Le Ven, A., Villy, M.C., Silveira, A.B., Houy, A., Masliah-Planchon, J., Warcoin, M., Le Mentec, M., Simaga, F., De Pauw, A., Buecher, B. et al. (2025) Uveal Melanoma and the Lynch Syndrome Tumor Spectrum. JAMA Ophthalmol, 143, 661–668.

21. Palles, C., West, H.D., Chew, E., Galavotti, S., Flensburg, C., Grolleman, J.E., Jansen, E.A.M., Curley, H., Chegwidden, L., Arbe-Barnes, E.H. et al. (2022) Germline MBD4 deficiency causes a multi-tumor predisposition syndrome. American Journal of Human Genetics, 109, 953–960.

22. Villy, M.C., Le Ven, A., Le Mentec, M., Masliah-Planchon, J., Houy, A., Bieche, I., Vacher, S., Vincent-Salomon, A., Dubois d’Enghien, C., Schwartz, M. et al. (2024) Familial uveal melanoma and other tumors in 25 families with monoallelic germline MBD4 variants. J. Natl. Cancer Inst., 116, 580–587.

23. Farlik, M., Sheffield, N.C., Nuzzo, A., Datlinger, P., Schonegger, A., Klughammer, J. and Bock, C. (2015) Single-cell DNA methylome sequencing and bioinformatic inference of epigenomic cell-state dynamics. Cell Rep, 10, 1386–1397.

24. Zuo, E., Sun, Y., Yuan, T., He, B., Zhou, C., Ying, W., Liu, J., Wei, W., Zeng, R., Li, Y. et al. (2020) A rationally engineered cytosine base editor retains high on-target activity while reducing both DNA and RNA off-target effects. Nat Methods, 17, 600-604.

25. Komor, A.C., Kim, Y.B., Packer, M.S., Zuris, J.A. and Liu, D.R. (2016) Programmable editing of a target base in genomic DNA without double-stranded DNA cleavage. Nature, 533, 420–424.

26. Liu, Z., Lu, Z., Yang, G., Huang, S., Li, G., Feng, S., Liu, Y., Li, J., Yu, W., Zhang, Y. et al. (2018) Efficient generation of mouse models of human diseases via ABE- and BE-mediated base editing. Nat Commun, 9, 2338.

27. Zou, X., Koh, G.C.C., Nanda, A.S., Degasperi, A., Urgo, K., Roumeliotis, T.I., Agu, C.A., Badja, C., Momen, S., Young, J. et al. (2021) A systematic CRISPR screen defines mutational mechanisms underpinning signatures caused by replication errors and endogenous DNA damage. Nat Cancer, 2, 643–657.

28. Degasperi, A., Zou, X., Amarante, T.D., Martinez-Martinez, A., Koh, G.C.C., Dias, J.M.L., Heskin, L., Chmelova, L., Rinaldi, G., Wang, V.Y.W. et al. (2022) Substitution mutational signatures in whole-genome-sequenced cancers in the UK population. Science, 376, abl9283.

29. Jumper, J., Evans, R., Pritzel, A., Green, T., Figurnov, M., Ronneberger, O., Tunyasuvunakool, K., Bates, R., Zidek, A., Potapenko, A. et al. (2021) Highly accurate protein structure prediction with AlphaFold. Nature, 596, 583–589.

30. Bret, H., Gao, J., Zea, D.J., Andreani, J. and Guerois, R. (2024) From interaction networks to interfaces, scanning intrinsically disordered regions using AlphaFold2. Nat Commun, 15, 597.

31. Steinegger, M. and Soding, J. (2017) MMseqs2 enables sensitive protein sequence searching for the analysis of massive data sets. Nat Biotechnol, 35, 1026–1028.

32. Steinegger, M., Meier, M., Mirdita, M., Vohringer, H., Haunsberger, S.J. and Soding, J. (2019) HH-suite3 for fast remote homology detection and deep protein annotation. BMC Bioinformatics, 20, 473.

33. Katoh, K. and Standley, D.M. (2013) MAFFT multiple sequence alignment software version 7: improvements in performance and usability. Mol Biol Evol, 30, 772–780.

34. Mirdita, M., Schutze, K., Moriwaki, Y., Heo, L., Ovchinnikov, S. and Steinegger, M. (2022) ColabFold: making protein folding accessible to all. Nat Methods, 19, 679–682.

35. Evans, R., O’Neill, M., Pritzel, A., Antropova, N., Senior, A., Green, T., Žídek, A., Bates, R., Blackwell, S., Yim, J. et al. (2021) Protein complex prediction with AlphaFold-Multimer. bioRxiv, 2021.2010.2004.463034.

36. Varga, J.K., Ovchinnikov, S. and Schueler-Furman, O. (2025) actifpTM: a refined confidence metric of AlphaFold2 predictions involving flexible regions. Bioinformatics, 41.

37. Pettersen, E.F., Goddard, T.D., Huang, C.C., Meng, E.C., Couch, G.S., Croll, T.I., Morris, J.H. and Ferrin, T.E. (2021) UCSF ChimeraX: Structure visualization for researchers, educators, and developers. Protein Sci, 30, 70–82.

38. Waterhouse, A.M., Procter, J.B., Martin, D.M., Clamp, M. and Barton, G.J. (2009) Jalview Version 2--a multiple sequence alignment editor and analysis workbench. Bioinformatics, 25, 1189-1191.

39. Rees, H.A. and Liu, D.R. (2018) Base editing: precision chemistry on the genome and transcriptome of living cells. Nat Rev Genet, 19, 770–788.

40. Trotter, E.W. and Hagan, I.M. (2020) Release from cell cycle arrest with Cdk4/6 inhibitors generates highly synchronized cell cycle progression in human cell culture. Open Biol, 10, 200200.

41. Savva, R., McAuley-Hecht, K., Brown, T. and Pearl, L. (1995) The structural basis of specific base-excision repair by uracil-DNA glycosylase. Nature, 373, 487–493.

42. Krokan, H.E., Drablos, F. and Slupphaug, G. (2002) Uracil in DNA--occurrence, consequences and repair. Oncogene, 21, 8935–8948.

43. Sanders, M.A., Chew, E., Flensburg, C., Zeilemaker, A., Miller, S.E., Al Hinai, A.S., Bajel, A., Luiken, B., Rijken, M., McLennan, T. et al. (2018) MBD4 guards against methylation damage and germ line deficiency predisposes to clonal hematopoiesis and early-onset AML. Blood, 132, 1526–1534.

44. Baljinnyam, T., Sowers, M.L., Hsu, C.W., Conrad, J.W., Herring, J.L., Hackfeld, L.C. and Sowers, L.C. (2022) Chemical and enzymatic modifications of 5-methylcytosine at the intersection of DNA damage, repair, and epigenetic reprogramming. PLoS One, 17, e0273509.

45. Orndorff, P.B., Poddar, S., Owens, A.M., Kumari, N., Ugaz, B.T., Amin, S., Van Horn, W.D., van der Vaart, A. and Levitus, M. (2023) Uracil-DNA glycosylase efficiency is modulated by substrate rigidity. Sci Rep, 13, 3915.

46. Kuijk, E., Jager, M., van der Roest, B., Locati, M.D., Van Hoeck, A., Korzelius, J., Janssen, R., Besselink, N., Boymans, S., van Boxtel, R. et al. (2020) The mutational impact of culturing human pluripotent and adult stem cells. Nat Commun, 11, 2493.

47. Koh, G.C.C., Nanda, A.S., Rinaldi, G., Boushaki, S., Degasperi, A., Badja, C., Pregnall, A.M., Zhao, S.J., Chmelova, L., Black, D. et al. (2025) A redefined InDel taxonomy provides insights into mutational signatures. Nat Genet, 57, 1132–1141.

48. Hamad, R.S. and Ibrahim, M.E. (2022) CMMRD caused by PMS1 mutation in a sudanese consanguineous family. Hered Cancer Clin Pract, 20, 16.

49. Alghamdi, B., Al-Hindi, H., Murugan, A.K. and Alzahrani, A.S. (2023) Thyroid Cancer, Neuroendocrine Tumor, Adrenal Adenoma, and Other Tumors in a Patient With a Germline PMS1 Mutation. J Endocr Soc, 7, bvad035.

50. Caulfield, M., Davies, J., Dennys, M., Elbahy, L., Fowler, T., Hill, S.J. and al., e. (2020).

51. Johansson, P.A., Brooks, K., Newell, F., Palmer, J.M., Wilmott, J.S., Pritchard, A.L., Broit, N., Wood, S., Carlino, M.S., Leonard, C. et al. (2020) Whole genome landscapes of uveal melanoma show an ultraviolet radiation signature in iris tumours. Nat Commun, 11, 2408.

52. Haradhvala, N.J., Kim, J., Maruvka, Y.E., Polak, P., Rosebrock, D., Livitz, D., Hess, J.M., Leshchiner, I., Kamburov, A., Mouw, K.W. et al. (2018) Distinct mutational signatures characterize concurrent loss of polymerase proofreading and mismatch repair. Nat Commun, 9, 1746.

53. Meng, H., Harrison, D.J. and Meehan, R.R. (2015) MBD4 interacts with and recruits USP7 to heterochromatic foci. J Cell Biochem, 116, 476–485.

54. Mac Partlin, M., Homer, E., Robinson, H., McCormick, C.J., Crouch, D.H., Durant, S.T., Matheson, E.C., Hall, A.G., Gillespie, D.A. and Brown, R. (2003) Interactions of the DNA mismatch repair proteins MLH1 and MSH2 with c-MYC and MAX. Oncogene, 22, 819–825.

55. Morera, S., Grin, I., Vigouroux, A., Couve, S., Henriot, V., Saparbaev, M. and Ishchenko, A.A. (2012) Biochemical and structural characterization of the glycosylase domain of MBD4 bound to thymine and 5-hydroxymethyuracil-containing DNA. Nucleic Acids Res, 40, 9917–9926.

56. Pidugu, L.S., Bright, H., Lin, W.J., Majumdar, C., Van Ostrand, R.P., David, S.S., Pozharski, E. and Drohat, A.C. (2021) Structural Insights into the Mechanism of Base Excision by MBD4. J Mol Biol, 433, 167097.

57. Jiricny, J. (2006) MutLalpha: at the cutting edge of mismatch repair. Cell, 126, 239–241.

58. Bergeron, F., Auvre, F., Radicella, J.P. and Ravanat, J.L. (2010) HO* radicals induce an unexpected high proportion of tandem base lesions refractory to repair by DNA glycosylases. Proc Natl Acad Sci U S A, 107, 5528–5533.

59. Miller, C.J., Kim, G.Y., Zhao, X. and Usdin, K. (2020) All three mammalian MutL complexes are required for repeat expansion in a mouse cell model of the Fragile X-related disorders. PLoS Genet, 16, e1008902.

60. McLean, Z.L., Gao, D., Correia, K., Roy, J.C.L., Shibata, S., Farnum, I.N., Valdepenas-Mellor, Z., Kovalenko, M., Rapuru, M., Morini, E. et al. (2024) Splice modulators target PMS1 to reduce somatic expansion of the Huntington’s disease-associated CAG repeat. Nat Commun, 15, 3182.

61. Reyes, G.X., Zhao, B., Schmidt, T.T., Gries, K., Kloor, M. and Hombauer, H. (2020) Identification of MLH2/hPMS1 dominant mutations that prevent DNA mismatch repair function. Commun Biol, 3, 751.

62. Raschle, M., Marra, G., Nystrom-Lahti, M., Schar, P. and Jiricny, J. (1999) Identification of hMutLbeta, a heterodimer of hMLH1 and hPMS1. J Biol Chem, 274, 32368–32375.

